# D-amino acid-mediated translation arrest is modulated by the identity of the incoming aminoacyl-tRNA

**DOI:** 10.1101/323295

**Authors:** Rachel C. Fleisher, Virginia W. Cornish, Ruben L. Gonzalez

## Abstract

A complete understanding of the determinants that restrict D-amino acid incorporation by the ribosome, which is of interest to both basic biologists as well as the protein engineering community, remains elusive. Previously, we demonstrated that D-amino acids are successfully incorporated into the C-terminus of the nascent polypeptide chain. Ribosomes carrying the resulting peptidyl-D-aminoacyl-tRNA (peptidyl-D-aa-tRNA) donor substrate, however, partition into subpopulations that either undergo translation arrest through inactivation of the ribosomal peptidyl-transferase center (PTC) or remain translationally competent. The proportion of each subpopulation is determined by the identity of the D-amino acid sidechain. Here, we demonstrate that the identity of the aminoacyl-tRNA (aa-tRNA) acceptor substrate that is delivered to ribosomes carrying a peptidyl-D-aa-tRNA donor further modulates this partitioning. Our discovery demonstrates that it is the pairing of the peptidyl-D-aa-tRNA donor and the aa-tRNA acceptor that determines the activity of the PTC. Moreover, we provide evidence that both the amino acid and tRNA components of the aa-tRNA donor contribute synergistically to the extent of arrest. The results of this work deepen our understanding of the mechanism of D-amino acid-mediated translation arrest and how cells avoid this precarious obstacle, reveal similarities to other translation arrest mechanisms involving the PTC, and provide a new route for improving the yields of engineered proteins containing D-amino acids.

---

Peptide bond formation is catalyzed by the ribosomal peptidyl-transferase center (PTC). This reaction involves transfer of the nascent polypeptide chain from the peptidyl-tRNA donor in the ribosomal peptidyl-tRNA binding (P) site to the aminoacyl-tRNA (aa-tRNA) acceptor in the ribosomal aa-tRNA binding (A) site (Fig. 1A). Although the PTC has evolved to catalyze peptide bond formation with a remarkable diversity of proteinogenic amino acids, these substrates are exclusively L enantiomers or achiral. Nonetheless, D-amino acids can be found in the cell at high concentrations^1^. Moreover, it is well-established that at least some aa-tRNA synthetase enzymes are capable of aminoacylating D-amino acids onto tRNAs^2,3^, suggesting the possibility that the translation machinery (TM) may have to contend with D-aa-tRNAs *in vivo*. Consistent with this possibility, a D-aa-tRNA deacylase enzyme has been identified that is nearly universally conserved and serves to remove D-amino acids from the tRNAs onto which they have been misacylated^4^. Although these observations suggest that D-aa-tRNAs might present a challenge to the TM, how D-aa-tRNAs that are successfully delivered to the translating ribosome impact protein synthesis remains poorly understood, with reported D-amino acid incorporation efficiencies ranging anywhere from zero to 40% or higher for incorporation of a single D-amino acid^3,5–7^. Nonetheless, continued efforts to develop a complete mechanistic understanding of how D-aa-tRNAs impede translation and/or how the TM might be engineered to facilitate the incorporation of D-aa-tRNAs are important for elucidating the limits of basic biology and enabling new directions in protein engineering^5,8^.

**Figure 1.**
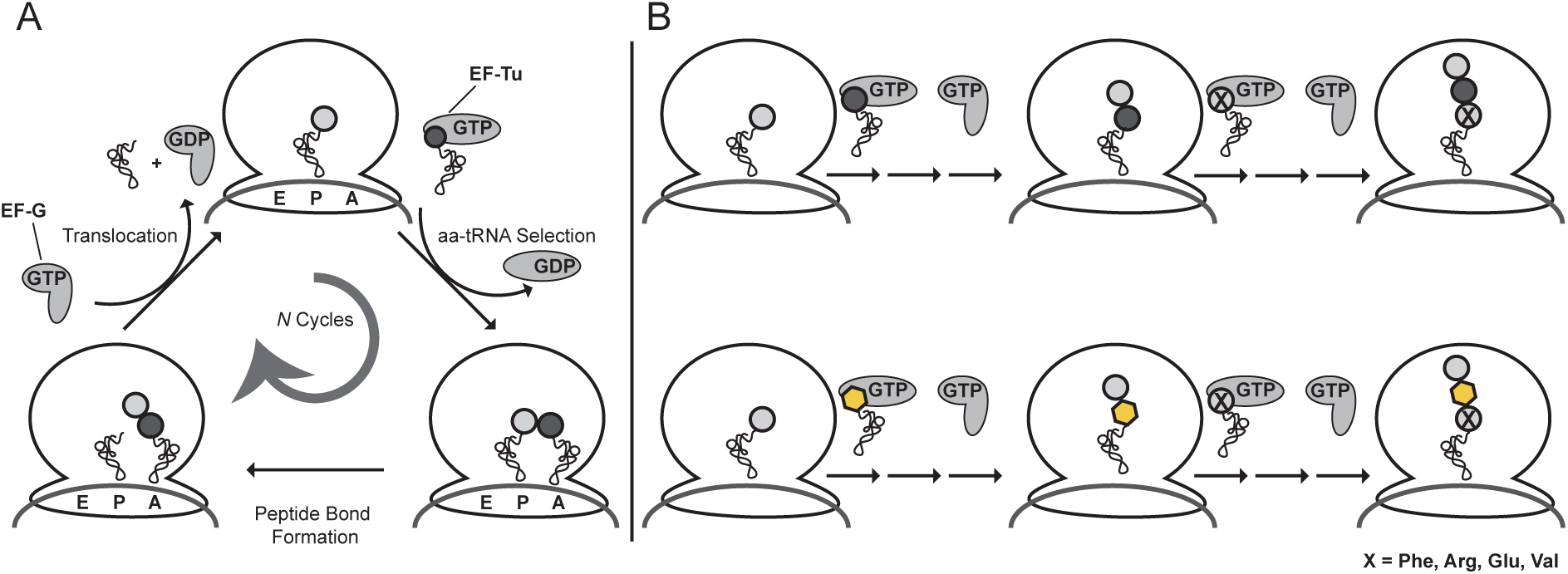
The experimental framework used to examine the role of the incoming aa-tRNA acceptor in D-amino acid-mediated translation arrest. (A) The translation elongation cycle begins with delivery of EF-Tu(GTP)aa-tRNA complexes to an EC carrying a peptidyl-tRNA donor at the P site. Following selection of the correct, mRNA-encoded EF-Tu(GTP)aa-tRNA, GTP is hydrolyzed, EF-Tu(GDP) dissociates from the aa-tRNA and the EC, and the aa-tRNA is accommodated into the A site. Upon accommodation, peptide bond formation results in transfer of the nascent polypeptide chain from the peptidyl-tRNA donor at the P site to the aa-tRNA acceptor at the A site. Subsequently, EF-G(GTP) catalyzes translocation of the P- and A-site tRNAs into the ribosomal tRNA exit (E) and P sites, with concomitant movement of the mRNA by one codon. (B) Tripeptide synthesis reactions were performed by delivering a mixture of either EF-Tu(GTP)L-Lys-tRNA^Lys^ (top row) or EF-Tu(GTP)D-Lys-tRNA^Lys^ (bottom row), EF-Tu(GTP)X-tRNA^X^, and EF-G(GTP) to ribosomal initiation complexes programmed with an mRNA encoding an fMet-Lys-X tripeptide (Table S1) and carrying a P-site f-[^35^S]-Met-tRNA^fMet^ (Supporting Materials and Methods).

Using a highly purified, *Escherichia coli*-based, *in vitro* translation system devoid of D-aa-tRNA deacylase and a series of biochemical assays reporting on the major steps of the translation elongation cycle (Fig. 1A), we recently published a mechanistic study of D-amino acid incorporation by the TM^9^. The results of this study demonstrated that D-aa-tRNA acceptors in complex with the GTP bound-form of elongation factor (EF)-Tu (*i.e.*, EF-Tu(GTP)D-aa-tRNA complexes) can be delivered to the A site and participate in peptide bond formation with yields mirroring those obtained from delivery of the corresponding L-aa-tRNA acceptors. Nonetheless, EF-G(GTP)-catalyzed translocation of the resulting peptidyl-D-aa-tRNA from the A- to P sites resulted in ribosomal elongation complexes (ECs) that partition into two subpopulations. One subpopulation is unable to undergo peptidyl transfer with the next aa-tRNA acceptor and becomes translationally arrested. The second subpopulation can undergo peptidyl transfer with the next aa-tRNA acceptor and remains translationally competent. Remarkably, the fraction of ECs that undergo translation arrest or remain translationally competent is dependent on the side chain of the D-amino acid, suggesting that the observed arrest arises from a D-amino acid side chain-dependent stabilization of an inactive conformation of the PTC. This conclusion was supported by chemical probing experiments and molecular dynamics simulations indicating that the peptidyl-D-aa-tRNA donor stabilizes conformations of 23S ribosomal RNA nucleotides A2058, A2059, A2062, A2063, G2505, and U2506 that are different from those which the corresponding peptidyl-L-aa-tRNA donor stabilizes. These nucleotides comprise part of the ribosomal nascent polypeptide exit tunnel entrance (ETE), a region of the ribosome that evidence suggests is conformationally and functionally coupled to the PTC^10^.

A striking feature of the study described above is that several of the nucleotides we identified are also implicated in macrolide^11^-, chloramphenicol^12^-, and nascent polypeptide-mediated translation arrest mechanisms^13,14^. Perhaps most interestingly, a growing number of studies are finding that the extent of arrest in these mechanisms is modulated by the identity of the peptidyl-aa-tRNA donor and aa-tRNA acceptor pair in the P- and A sites of the arrested EC^15–^17. Motivated by these studies, we hypothesized that such a ‘pairing effect’ may likewise modulate the extent of D-amino acid-mediated translation arrest. To test this hypothesis, we have investigated whether and how the identity of the aa-tRNA acceptor can modulate the fraction of ECs carrying a peptidyl-D-aa-tRNA donor that undergo translation arrest or remain translationally competent.

Using our previously described *in vitro* translation system^9^, we began our investigation by performing fMet-L-Lys-X and fMet-D-Lys-X tripeptide synthesis reactions in which the identity of the third amino acid, denoted by the ‘X’, was varied by delivering EF-Tu(GTP)aa-tRNAs formed using one of four different aa-tRNA acceptors (Supporting Materials and Methods and Figs. S1 and 1B). The choice of a peptidyl-D-Lys-tRNA^Lys^ donor in these experiments was driven by our previous demonstration that the fraction of ECs that undergo translation arrest is closest to 50 % when a peptidyl-D-Lys-tRNA^Lys^ donor is paired with a Phe-tRNA^Phe^ acceptor^9^. We therefore reasoned that using the peptidyl-D-Lys-tRNA^Lys^ donor and Phe-tRNA^Phe^ acceptor pairing as a reference would allow us to most easily detect both increases and decreases in the fraction of ECs that undergo translation arrest. For the aa-tRNA acceptors, we chose aa-tRNAs in which the amino acids differed by hydrophobicity (Phe-tRNA^Phe^), electrostatics (Arg-tRNA^Arg^ and Glu-tRNA^Glu^), and beta-branching (Val-tRNA^Val^). To validate our experimental system, we performed an initial pair of tripeptide synthesis reactions using Phe-tRNA^Phe^ as the aa-tRNA acceptor (Figs. S2 and 2). The fMet-L-Lys-Phe and fMet-D-Lys-Phe tripeptide synthesis reactions exhibited end-point yields of 89 ± 1 % and 64 ± 1 %, respectively, in excellent agreement with our previous observations of 80 ± 1 % and 58 ± 5 %, respectively^9^.

**Figure 2.**
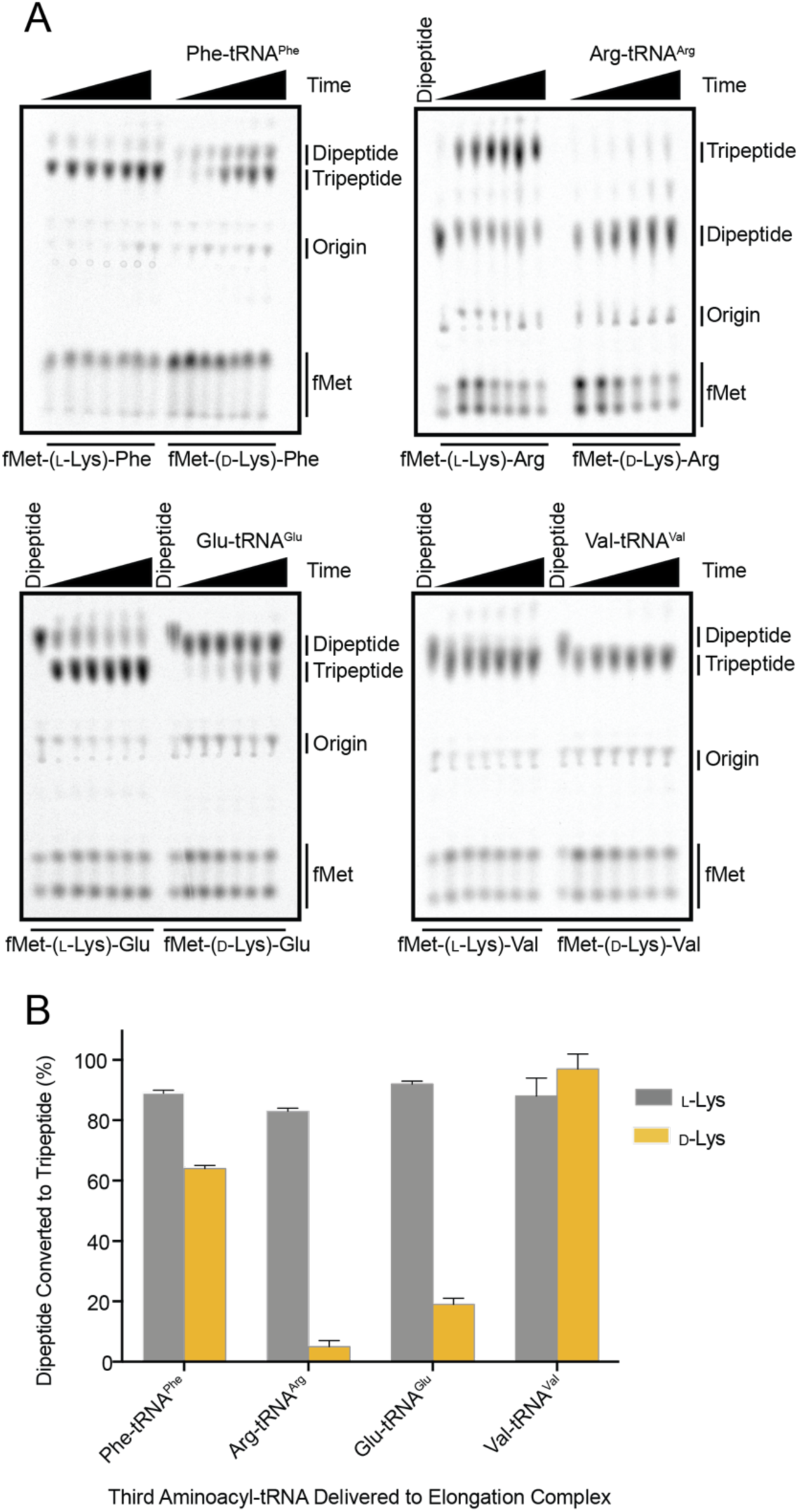
The identity of the incoming aa-tRNA acceptor modulates the extent of D-amino acid-mediated translation arrest. (A) Tripeptide synthesis reaction products were collected at reaction time points spanning 0–60 min in order to ensure that the reactions had gone to completion and we were measuring true end-point yields. Products were separated using electrophoretic thin layer chromatography (eTLC) and visualized by phosphorimaging (Supporting Materials and Methods). (B) End-point yields of tripeptide synthesis reactions. Reaction time courses were performed in duplicate and in all cases were observed to go to completion by 30 min (Fig. S2). Consequently, the duplicate yield measurements collected at the 30 and 60 min time points for each tripeptide synthesis reaction were clustered, averaged, and reported as the mean reaction end-point. Error bars represent the standard deviation of the mean (Supporting Materials and Methods).

Subsequently, we performed tripeptide synthesis reactions using Arg-tRNA^Arg^ as the aa-tRNA acceptor (Figs. S2 and 2). In this case, the fMet-L-Lys-Arg and fMet-D-Lys-Arg tripeptide synthesis reactions exhibited end-point yields of 83 ± 1 % and 5 ± 2 %, respectively. The latter result stands in stark contrast to the end-point yield of 64 ± 1 % observed in the fMet-D-Lys-Phe tripeptide synthesis reaction, revealing that the fraction of ECs that undergo D-amino acid-mediated translation arrest depends on the identity of the aa-tRNA acceptor.

Given the striking effect that we observed when the aa-tRNA acceptor carried a positively charged amino acid, we used the next tripeptide synthesis reactions to test the performance of an aa-tRNA acceptor carrying a negatively charged amino acid, Glu-tRNA^Glu^ (Figs. S2 and 2). Here, we observed end-point yields of 92 ± 1 % and 19 ± 2 % for the synthesis of the fMet-L-Lys-Glu and fMet-D-Lys-Glu tripeptides, respectively. The data we have obtained here using aa-tRNA acceptors carrying charged amino acids are notably similar to our previous findings in which delivery of a Lys-tRNA^Lys^ acceptor to an EC carrying either fMet-D-Phe-tRNA^Phe^ or fMet-D-Val-tRNA^Val^ donors results in 18 ± 3 % and 11 ± 1 % of the ECs undergoing translation arrest, respectively^9^. Collectively, our results strongly suggest that charged aa-tRNA acceptors exacerbate D-amino acid mediated translation arrest.

We next performed tripeptide synthesis reactions in which Val-tRNA^Val^ served as the incoming aa-tRNA (Figs. S2 and 2). Surprisingly, the fMet-L-Lys-Val and fMet-D-Lys-Val tripeptide synthesis reactions exhibited end-point yields of 88 ± 6 % and 97 ± 5 %, respectively. This result again highlights the ability of the aa-tRNA acceptor to modulate the extent of D-amino acid-mediated translation arrest. Perhaps most interestingly, these findings demonstrate that, by judiciously choosing the identity of the aa-tRNA acceptor, it is possible to completely bypass D-amino acid-mediated translation arrest and incorporate a D-amino acid into a ribosome-synthesized protein or polypeptide as effectively as the corresponding L-amino acid can be incorporated.

The ability of the aa-tRNA acceptor to modulate the extent of D-amino acid-mediated translation arrest can originate from either its amino acid- or tRNA component. In order to dissect the contributions of these components, we misacylated phenylalanine onto tRNA^Glu^ to generate Phe-tRNA^Glu^ (Supporting Materials and Methods and Fig. S1)^9,18^ and used it as the aa-tRNA acceptor in the next tripeptide synthesis reactions (Figs. S2 and 3). If the phenylalanine component is the major contributor, then we would expect an fMet-D-Lys-Phe tripeptide end-point yield that is similar or identical to that obtained upon delivery of Phe-tRNA^Phe^ (*i.e.*, 64 ± 1 %). In contrast, if the tRNA^Glu^ component is the major contributor, then we would expect an fMet-D-Lys-Phe tripeptide end-point yield that is similar or identical to that obtained upon delivery of Glu-tRNA^Glu^ (*i.e.*, 19 ± 2 %). Contrary to either expectation, fMet-L-Lys-Phe and fMet-D-Lys-Phe tripeptide synthesis reactions using Phe-tRNA^Glu^ as the aa-tRNA acceptor exhibited end-point yields of 95 ± 3 % and 95 ± 2 %, respectively. These results demonstrate that both the amino acid and tRNA^19^ components of the aa-tRNA acceptor contribute to modulating the extent of D-amino acid-mediated translation arrest and do so synergistically.

**Figure 3.**
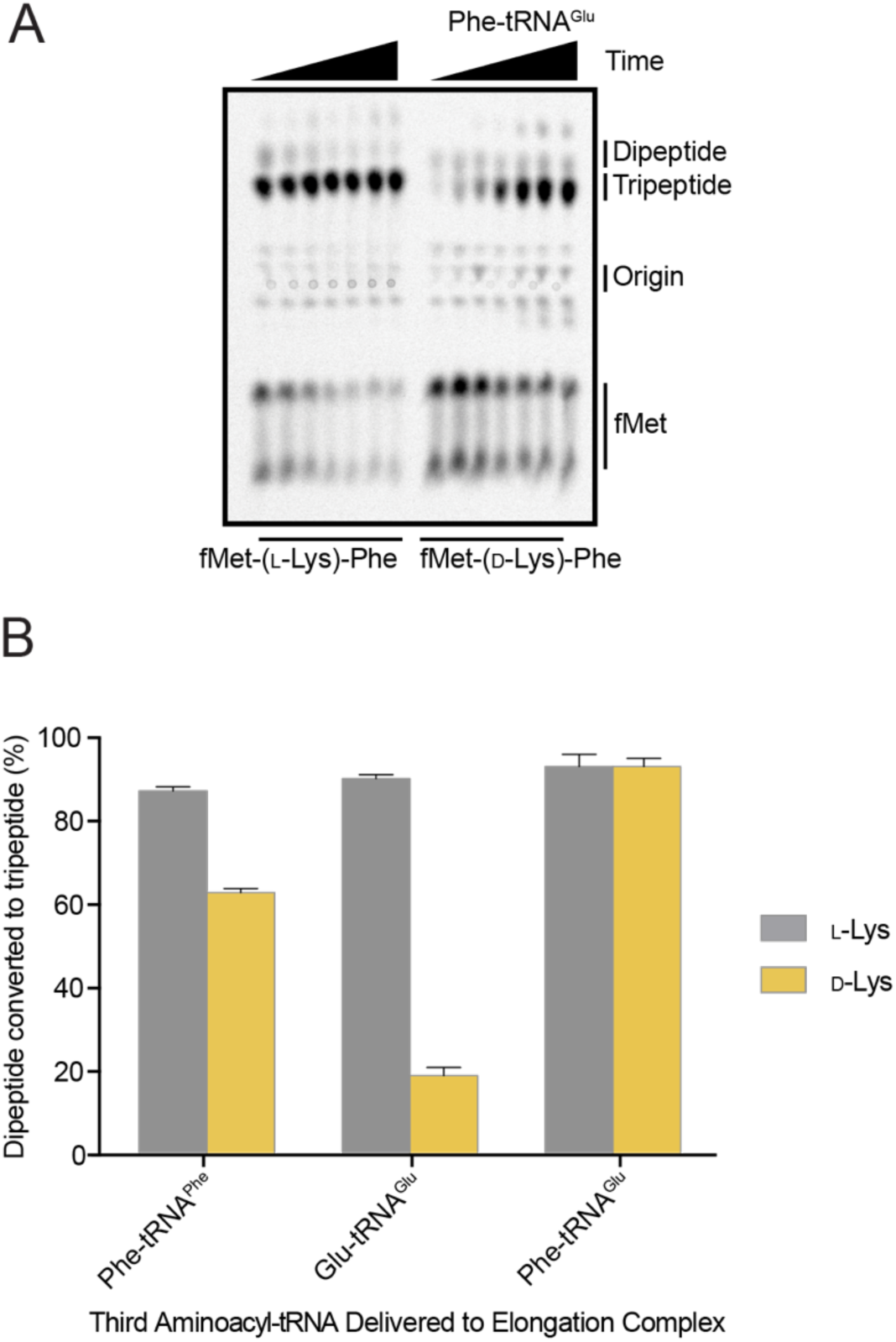
The amino acid- and tRNA components of the aa-tRNA acceptor contribute synergistically to the extent of D-amino acid-mediated translation arrest. (A) eTLC of the fMet-L-Lys-Phe and fMet-D-Lys-Phe tripeptide synthesis reactions performed using Phe-tRNA^Glu^ as the aa-tRNA acceptor. (B) End-point yields of tripeptide synthesis reactions. Reaction time courses were performed in duplicate and in all cases were observed to go to completion by 30 min (Fig. S2). Consequently, the duplicate yield measurements collected at the 30 and 60 min time points for each tripeptide synthesis reaction were clustered, averaged, and reported as the mean reaction end-point. Error bars represent the standard deviation of the mean (Supporting Materials and Methods).

In this communication, we demonstrate that the identity of the incoming aa-tRNA acceptor can modulate the fraction of ECs carrying a peptidyl-D-aa-tRNA donor that undergo D-amino acid-mediated translation arrest. In addition, we show that the amino acid- and tRNA components of the incoming aa-tRNA contribute synergistically to modulating the extent of translation arrest. Extending our previously proposed model for D-amino acid-mediated translation arrest^9^, our data strongly suggests that translocation of the peptidyl-D-aa-tRNA into the P site alters the conformational dynamics of the PTC. Specifically, the peptidyl-D-aa-tRNA likely limits the conformational space that the PTC can sample and, to an extent that depends on the identity of the D-amino acid side chain, predisposes the PTC to stably adopt an inactive conformation. The incoming aa-tRNA acceptor can then further modulate the dynamics of the PTC, biasing the PTC towards stably adopting an inactive conformation that induces translation arrest or an active conformation that enables continued translation. The ability of a particular aa-tRNA acceptor to bias the PTC towards stably adopting an inactive conformation establishes the fraction of ECs that will undergo translation arrest. Thus, the extent of D-amino acid-mediated translation arrest is not determined solely by the side chain of the D-amino acid, as we originally proposed, but rather by the identity of a particular peptidyl-D-aa-tRNA donor and aa-tRNA acceptor pair (Fig. 4).

**Figure 4.**
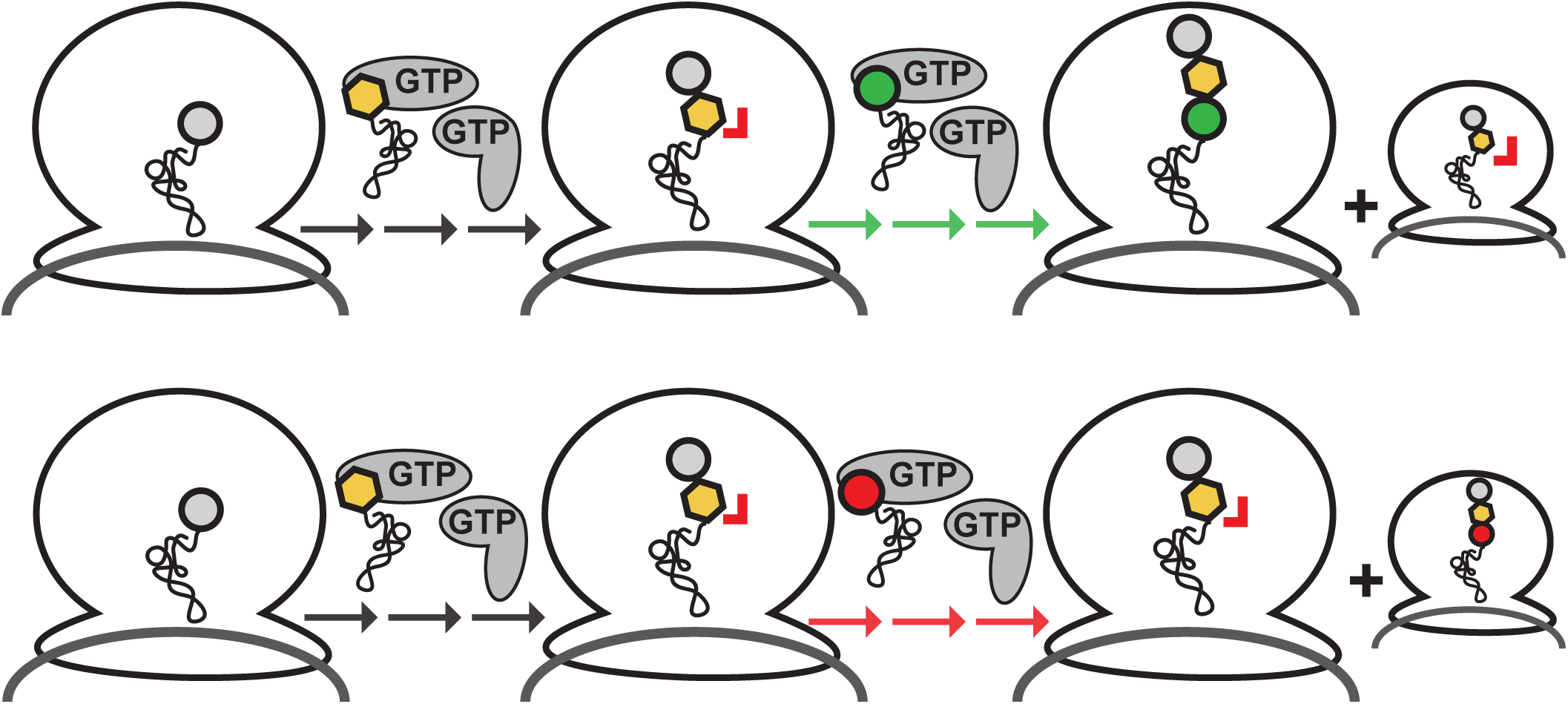
Extended mechanistic model for D-amino acid-mediated translation arrest. Translocation of a peptidyl-D-aa-tRNA donor into the P site predisposes the PTC to stably adopt an inactive conformation. Pairing of a particular peptidyl-D-aa-tRNA donor with an aa-tRNA acceptor that facilitates the ability of the PTC to stably adopt an inactive conformation results in a large fraction of ECs undergoing translation arrest (red aa-tRNA acceptor and pathway), whereas pairing with an aa-tRNA acceptor that interferes with the ability of the PTC to stably adopt an inactive conformation results in a large fraction of ECs remaining translationally competent (green aa-tRNA acceptor and pathway).

Previously, we have demonstrated that translocation of a peptidyl-D-aa-tRNA into the P site allosterically induces a conformational change of 23S rRNA nucleotides that comprise the ETE. Intriguingly, several other translation arrest mechanisms involving these same ETE nucleotides likewise demonstrate a pairing effect similar to that described here. For example, in their studies of a nascent polypeptide-mediated translation arrest mechanism involving contacts of the nascent polypeptide with these ETE nucleotides^20^, Mankin and co-workers demonstrated that the extent to which they could exacerbate or relieve the corresponding nascent polypeptide-mediated translation arrest depended on the identity of the aa-tRNA acceptor^16^. Surprisingly, they observed that Val-tRNA^Val^ and Phe-tRNA^Phe^ acceptors exhibited among the lowest arrest efficiencies, while Arg-tRNA^Arg^ and Glu-tRNA^Glu^ acceptors exhibited among the highest, a trend that is consistent with the trend that we have observed here. Mankin and co-workers have also shown that binding of a macrolide antibiotic to these ETE nucleotides robustly induces translation arrest primarily in the context of specific, problematic peptidyl-aa-tRNA donor and aa-tRNA acceptor pairs^15^. Similar conformational and pairing effects were observed in the context of the PTC-binding antibiotics chloramphenicol and linezolid^11,17^. Thus, our findings contribute to an emerging understanding of a conformational and functional link between the ETE and the PTC, in which translation arrest can be induced by particular peptidyl-aa-tRNA donor and aa-tRNA acceptor pairs. In addition, the results we report here demonstrate the potentially devastating effects that incorporation of a D-amino acid into the nascent polypeptide chain can have on translation and highlight the important role of that D-aa-tRNA deacylase enzymes play in protecting ECs under stress conditions in which D-amino acids may be misacylated onto tRNAs and delivered to ECs^21^. Finally, our results reveal a straightforward approach for improving the efficiency with which D-amino acids can be incorporated into proteins or polypeptides for protein engineering applications: By simply optimizing the pairing of peptidyl-D-aa-tRNA donors and aa-tRNA acceptors, the efficiency of incorporation can be improved dramatically without the need for complex engineering of the TM.

## Acknowledgements

We thank Dr. Colin Kinz-Thompson for his assistance in data analysis. R.C.F was supported by the Training Program in Molecular Biophysics at Columbia University (T32 GM008281). This work was supported in part by funds to V.W.C. and R.L.G. from Columbia University as well as National Institutes of Health (NIH) Grant R01 GM090126 (to R.L.G. and V.W.C.) and R01 GM119386 (to R.L.G.).

## Author Contributions

R.C.F. and R.L.G. designed the research; R.C.F. performed the experiments and analyzed the data; R.C.F. and R.L.G. interpreted the results. R.C.F. and R.L.G. wrote the manuscript; all three authors approved the final manuscript.

